# A TLR2-associated entry pathway promotes productive dengue virus serotype 2 infection and inflammatory responses in classical monocytes

**DOI:** 10.64898/2026.09.30.755835

**Authors:** Jorge Andrés Castillo, Madhura Punekar, Vinit Upasani, Peter van Duinkerken, Bram ter Ellen, Heidi van der Ende-Metselaar, Silvio Urcuqui-Inchima, Jolanda M. Smit, Izabela A. Rodenhuis-Zybert

## Abstract

Severe dengue is characterized by systemic inflammation and vascular leakage. Recent evidence indicates that Toll-like receptor 2 (TLR2) supports early steps of dengue virus (DENV) infection in monocytes, whereas intracellular viral RNA sensing drives much of the subsequent inflammatory response. However, whether this entry-associated function of TLR2 differs among human monocyte subsets remains unknown. Here, we investigated how the TLR2 axis regulates DENV serotype 2 (DENV2) infection and inflammatory output in purified primary human monocytes and sorted monocyte subsets. Blocking TLR2 at the cell surface or targeting its intracellular TIR domain reduced virus-cell association, productive infection, and the release of inflammatory and antiviral mediators. Among TLR2-associated receptors, CD14 and, to a lesser extent, CD36 contributed to efficient infection, whereas TLR1, TLR6, and TLR10 were dispensable under the conditions tested. Notably, TLR2 blockade reduced viral production and cytokine release in CD16^−^ classical monocytes but not in pooled CD16^+^ monocytes, identifying a subset-specific dependence on the TLR2 axis. TLR3 contributed to inflammatory and antiviral responses without affecting viral replication, consistent with post-entry recognition of viral RNA. Finally, blocking the TLR2 axis during monocyte infection prevented the loss of endothelial barrier integrity induced by monocyte-derived soluble mediators. Together, these findings identify a TLR2-associated entry route that preferentially supports productive DENV2 infection in classical monocytes and thereby enables RNA-sensor-dependent inflammatory responses that compromise endothelial function.

**AUTHOR SUMMARY:** Monocytes are major targets of dengue virus infection and can release inflammatory mediators that weaken the endothelial barrier, a hallmark of severe dengue. We recently showed that TLR2 acts early during direct and antibody-enhanced dengue virus infection, whereas intracellular receptors that detect viral RNA contribute to the subsequent inflammatory response. Here, we show that this pathway is not used equally by all monocytes. Classical CD16^−^ monocytes depended strongly on the TLR2 axis for productive infection and cytokine release, whereas pooled CD16^+^ monocytes did not. CD14 and, more modestly, CD36 contributed to infection in purified monocytes. Blocking TLR2 during infection also prevented monocyte-derived factors from disrupting endothelial barrier function. These findings suggest that monocyte identity determines how dengue virus enters cells and how strongly infection is coupled to potentially harmful inflammation.

## INTRODUCTION

The four dengue virus serotypes (DENV1-4) co-circulate in more than 120 countries and are estimated to cause 390 million infections annually, of which approximately 96 million are clinically apparent [1,2]. Most symptomatic infections are resolved after an acute febrile illness, but a subset progresses to severe dengue, characterized by thrombocytopenia, hemorrhage, increased vascular permeability, plasma leakage, and organ impairment [1,3]. Excessive or prolonged production of inflammatory mediators is thought to contribute to endothelial dysfunction and disease progression [4,5]. However, the cellular and molecular events that couple DENV infection to vascular pathology remain incompletely defined.

DENV is an enveloped, positive-sense single-stranded RNA virus of the genus Orthoflavivirus in the family Flaviviridae. Viral entry requires receptor-mediated endocytosis, followed by low-pH-dependent fusion of the viral and endosomal membranes and release of the genome into the cytoplasm [6,7]. Viral translation and replication subsequently occur in endoplasmic reticulum-derived membrane structures [8,9]. Although several cell types can support DENV infection in vitro, cells of the mononuclear phagocyte lineage, including dendritic cells, macrophages, and monocytes, are considered important targets in vivo [10–12]. DC-SIGN, the mannose receptor, heparan sulfate, and a CD14-associated factor have been implicated in DENV attachment or entry in dendritic cells and macrophages [13–15]. The host factors that determine susceptibility of circulating human monocytes remain less well defined.

Human monocytes comprise CD14^++^CD16^−^ classical monocytes (CM), CD14^++^CD16^+^ intermediate monocytes (IM), and CD14^+^CD16^++^ non-classical monocytes (NM) [16–18]. These subsets differ in abundance, differentiation state, receptor expression, trafficking, and effector function [17–19]. Acute dengue is associated with reduced frequencies of CM and expansion of CD16^+^ populations [20–22]. In our previous clinical study, CM frequency and sustained high TLR2 expression specifically on CM were associated with progression to severe disease [20]. These observations suggested that the biological consequences of TLR2 expression may differ among monocyte subsets.

Our initial work identified TLR2 and CD14 as determinants of DENV-induced responses in PBMCs [20]. More recent work refined this model. During direct and antibody-dependent enhancement (ADE) infection, TLR2 activity was required early, whereas UV-inactivated DENV2 did not trigger the measured inflammatory response and blockade of TLR2 late after infection was ineffective. TLR3 and TLR8 contributed to cytokine production without altering infection, supporting a serial model in which TLR2 licenses entry or early post-entry events and intracellular RNA sensors generate much of the downstream response [23]. This model raises an important unresolved question: does the TLR2-associated entry route operate equally across human monocyte subsets?

TLR2 function is shaped by associated receptors and membrane context. CD14 and CD36 can facilitate ligand transfer and internalization of TLR2-containing complexes, TLR1 and TLR6 form signaling heterodimers with TLR2, and TLR10 can modulate TLR2-dependent responses [24–27]. Because these molecules are differentially expressed among monocyte subsets, receptor context rather than TLR2 abundance alone may determine whether a monocyte uses a TLR2-associated route of infection.

Here, we tested the hypothesis that monocyte subsets differ in their dependence on a TLR2-associated entry pathway and that this difference determines the magnitude of downstream RNA-sensor-dependent inflammation. Using enriched primary monocytes, sorted CD16^−^ and pooled CD16^+^ monocytes, complementary TLR2-inhibition strategies, assays of virus-cell association and membrane fusion, and functional endothelial barrier models, we defined the subset-specific contribution of TLR2 and associated receptors to DENV2 infection and inflammatory output.

## MATERIALS AND METHODS

### Ethics statement

PBMCs were obtained from anonymized buffy coats from healthy blood donors through Sanquin Blood Bank. According to the principles expressed in the Declaration of Helsinki, written informed consent was obtained from all individuals who voluntarily donated blood to the Sanquin Blood Bank.

### PBMC isolation and monocyte enrichment or depletion

PBMCs were isolated from buffy coats using Ficoll-Paque density-gradient centrifugation, as described previously [28]. Cells were cryopreserved in FBS containing 10% DMSO. Monocytes were enriched by negative magnetic selection and subsequent plastic adherence. Cultures were used when purity exceeded 90%, as assessed by the absence of CD3^+^, CD19^+^, and CD56^+^ cells. For monocyte-depletion experiments, CD14^+^ cells were removed by positive magnetic selection. Monocyte-depleted PBMCs were used when depletion exceeded 80%.

### Sorting of monocyte populations

PBMCs were stained with antibodies against CD3, CD19, CD56, HLA-DR, CD14, and CD16. CD14^+^CD16^−^ classical monocytes and pooled CD14^+^CD16^+^ intermediate/non-classical monocytes were sorted using a Sony SH800S cell sorter. Purity exceeded 96%; recovery exceeded 85% for CD16^−^ cells and 70% for pooled CD16^+^ cells. Sorted cells were allowed to adhere for 2 h in RPMI-1640 containing 0.5% FBS before the medium was replaced with RPMI-1640 containing 10% FBS.

### Cells

C6/36 mosquito cells (ATCC) were maintained in MEM supplemented with 10% heat-inactivated fetal bovine serum (FBS), sodium bicarbonate, non-essential amino acids, 4 mM L-glutamine, penicillin, and streptomycin at 28°C with 5% CO_2_. BHK-21 clone 15 cells were maintained in DMEM containing 10% FBS, 4 mM L-glutamine, penicillin, and streptomycin at 37°C with 5% CO_2_. Primary human umbilical vein endothelial cells (HUVECs; Lonza) were cultured in EBM-2 supplemented with the EGM-2 SingleQuot kit. HEK-Blue hTLR2 cells (InvivoGen) were maintained according to the manufacturer’s instructions. BS-C-1 cells (ATCC CCL-26) were cultured in DMEM containing 10% FBS, 25 mM HEPES, penicillin, and streptomycin at 37°C with 5% CO_2_.

### Antibodies, agonists, and inhibitors

Antibodies used for immunophenotyping included reagents against CD3, CD14, CD16, CD19, CD36, CD56, HLA-DR, TLR1, TLR2, TLR6, TLR10, E-selectin, ICAM-1, and VCAM-1, together with the corresponding isotype controls. TLR2-axis components were blocked with antibodies against TLR2, TLR1, TLR6, TLR10, CD14, or CD36. The TLR2 TIR-domain inhibitor C29, NF-κB inhibitor Bay 11-7082, TLR8 inhibitor CU-CPT9a, and a TLR3/dsRNA-complex inhibitor were used as indicated. PAM3CSK4, PAM2CSK4, poly(I:C), and ssRNA40 were used as pathway controls. DENV2 infection was detected with flavivirus E-protein-specific monoclonal antibody 4G2 and fluorophore- or HRP-conjugated secondary antibodies.

### Virus production and quantification

DENV2 strain 16681 was propagated in C6/36 cells at a multiplicity of infection (MOI) of 0.05. Supernatants were collected seven days after infection, clarified at 800 × g for 10 min, aliquoted, and stored at −80°C. Infectious titers were determined by plaque assay in BHK-21 cells [29]. Viral genome-equivalent copies (GECs) were quantified by RT-qPCR using a plasmid encoding the full DENV2 genome as the standard [30].

### DENV2 infection and receptor or pathway inhibition

PBMCs, monocyte-depleted PBMCs, enriched monocytes, and sorted monocyte populations were exposed to DENV2 strain 16681 at an MOI of 10 in RPMI-1640 containing 2% FBS. After 2 h, the inoculum was removed, cells were washed, and infection was continued in RPMI-1640 containing 10% FBS for 48 h unless indicated otherwise. For inhibition experiments, cells were pretreated for 2 h with anti-TLR2 (7.5 µg/mL), anti-TLR1 (7.5 µg/mL), anti-TLR6 (7.5 µg/mL), anti-TLR10 (3 µg/mL), anti-CD14 (3 µg/mL), anti-CD36 (2.5 µg/mL), or the corresponding isotype control. C29 (100 nM), Bay 11-7082 (2 µM), CU-CPT9a (50 µM), and the TLR3 inhibitor (10 µM) were prepared in DMSO and compared with vehicle controls. Inhibitors were present during pretreatment, inoculation, and subsequent culture. Pathway activity and inhibitor performance were assessed using PAM3CSK4 (100 ng/mL), PAM2CSK4 (100 ng/mL), poly(I:C) (25 µg/mL), or ssRNA40 (1 µg/mL).

### Flow cytometry

DENV2 infection was quantified by intracellular staining for viral E protein. Cells were fixed in 4% paraformaldehyde, permeabilized with PBS containing 0.5% Tween-20, stained with 4G2, and subsequently stained with Alexa Fluor 647-conjugated anti-mouse IgG. Expression of TLR2-axis components was assessed in PBMC monocyte subsets using antibodies against TLR2, TLR1, TLR6, TLR10, CD36, CD14, CD16, and HLA-DR. HUVEC activation was quantified by surface expression of E-selectin, ICAM-1, and VCAM-1. Viability was assessed with fixable viability dye eFluor 780. Data were acquired on a BD FACSVerse and analyzed in FlowJo.

### Cytokine quantification

IL-1β, TNF-α, IL-6, IL-8, IL-10, IL-12p70, IP-10, GM-CSF, IFN-α2, IFN-β, IFN-λ1, IFN-λ2/3, and IFN-γ were quantified in cell-free supernatants using the LEGENDplex human anti-virus response panel. Data were acquired on a BD FACSVerse and analyzed with LEGENDplex software.

### Endothelial activation and permeability

Confluent HUVEC monolayers were stimulated for 6 h with cell-free supernatants from PBMC or monocyte cultures mixed 1:1 with endothelial medium. LPS (1 µg/mL) served as a positive control. Direct exposure of HUVECs to DENV2 was included to distinguish effects of the virus from effects of monocyte-derived soluble mediators. HUVEC activation was assessed by flow-cytometric measurement of E-selectin, ICAM-1, and VCAM-1.

For barrier assays, 5 × 10^4 HUVECs were seeded on collagen I-coated 0.4-µm transwell inserts and cultured until confluent. HUVECs were stimulated with monocyte supernatants diluted 1:1 in endothelial medium; TNF-α (100 ng/mL) served as a positive control. Transendothelial electrical resistance (TEER) was measured every 2 h until 12 h and again at 24 h. Relative TEER was calculated after subtracting resistance from cell-free inserts and normalized to untreated HUVECs. Dextran-FITC flux was measured by adding 200 µg/mL dextran-FITC to the upper chamber after 22 h and quantifying fluorescence in the lower chamber 2.5 h later.

### Cell-based virus-association assay

Poly-L-lysine-coated 96-well plates were seeded with 2 × 10^4 HEK-Blue hTLR2 cells per well. Cells were pretreated as indicated and exposed to DENV2 for 2 h. After removal of unbound virus, cells were fixed, permeabilized, and stained with 4G2 followed by HRP-conjugated anti-mouse IgG. Antibody complexes were detected with o-phenylenediamine dihydrochloride substrate, and absorbance was measured at 492 nm.

### DiD labeling and membrane-fusion assay

Purified DENV2 strain 16681 and CHIKV LR-2006 OPY1 strain were labeled with the lipophilic fluorescent probe DiD (1,10-dioctadecyl-3,3,30,30 -tetramethylindodicarbocyanine, 4-chlorobenzenesulfonate salt; Invitrogen #D7757), as described previously [31,32]. BS-C-1 cells (2 × 104 cells/well) were grown on µ-Slide 8 Well high chambered coverslip slides (Ibidi GmbH #80806) were pretreated for 1.5 h with C29 (100 µM), DMSO (vehicle control), and NH_4_Cl (50 mM) and exposed to DiD-labeled DENV2 for 30 min or CHIKV for 20 min at 37°C. Cells were washed twice with 1x PBS (with calcium and magnesium), fixed with 4% PFA, and imaged using a Leica THUNDER wide-field DMi8 microscope. 50-100 snapshots of randomly selected fields were acquired per condition. Fluorescent particle area in arbitrary units (AU) was quantified in ImageJ and averaged per experiment. NH_4_Cl was included as a control for inhibition of low-pH-dependent fusion.

### Statistical analysis

Statistical analyses were performed using GraphPad Prism version 10 (GraphPad Software, USA). The number of independent biological replicates is indicated in the corresponding figure legends; unless stated otherwise, each biological replicate represents an independent blood donor. Comparisons involving multiple treatment groups or experimental factors were performed using two-way ANOVA followed by Dunnett’s multiple-comparisons test, as specified in the figure legends. Selected datasets were analyzed using the Kruskal-Wallis test. Statistical significance was assessed at a 95% confidence level, with P < 0.05 considered statistically significant. Significance is indicated as *P < 0.05, **P < 0.01, and ***P < 0.001. The statistical test, sample size, normalization procedure, and reference control for each analysis are provided in the corresponding figure legend.

## RESULTS

### Monocytes are major sources of DENV2 replication and endothelial-activating mediators in PBMC cultures

Our previous studies identified monocytes as an important source of TLR2-dependent responses to DENV infection in PBMCs [20,23]. Before examining the underlying mechanism in purified cells, we first asked how strongly monocytes contributed to infection and inflammatory output in the mixed PBMC population. We therefore compared intact PBMCs with PBMCs depleted of CD14^+^ monocytes. Monocyte depletion substantially reduced the frequency of DENV2 E-positive cells, viral genome production, and the release of infectious progeny (Fig. S1A-C). The functional consequences were equally clear: supernatants from DENV2-infected PBMCs activated endothelial cells, whereas supernatants from monocyte-depleted cultures did not (Fig. S1D). These experiments established that, under our culture conditions, monocytes were major contributors to both productive DENV2 infection and the release of soluble endothelial-activating factors. We therefore continued with enriched monocyte cultures, consistently obtaining purities above 90% (Fig. S2).

### TLR2 activity supports early events required for productive DENV2 infection

Recent work from our group indicates that TLR2 acts primarily during the early phase of direct and antibody-enhanced DENV2 infection, rather than functioning as the sole sensor responsible for the subsequent cytokine response [23]. We therefore first examined whether TLR2 activity affected infection itself in purified primary monocytes. To reduce the likelihood that an effect was specific to a single reagent, we used two complementary approaches: an antibody directed against the extracellular domain of TLR2 and C29, a small-molecule inhibitor targeting its intracellular TIR domain [33]. Neither treatment reduced monocyte viability in mock-infected or DENV-infected cultures (Fig. S3A). As a functional control, both treatments markedly reduced the capacity of PAM3-stimulated monocyte supernatants to activate endothelial cells (Fig. S3B).

We next infected monocytes with DENV2 at an MOI of 10 and assessed infection 48 h later (Fig. 1A). Pretreatment with C29 reduced the infected cell mass by approximately 80%, whereas the TLR2-blocking antibody produced a reduction of approximately 50% relative to the corresponding controls (Fig. 1B; Fig. S4). The effect extended beyond the number of E-positive cells. C29 reduced viral genome and infectious-particle production by approximately 8- to 10-fold, while extracellular TLR2 blockade reduced both readouts by at least 100-fold (Fig. 1C, D). Thus, two mechanistically distinct forms of TLR2 inhibition converged on the same phenotype: inefficient productive infection of primary monocytes.

**Fig. 1.**
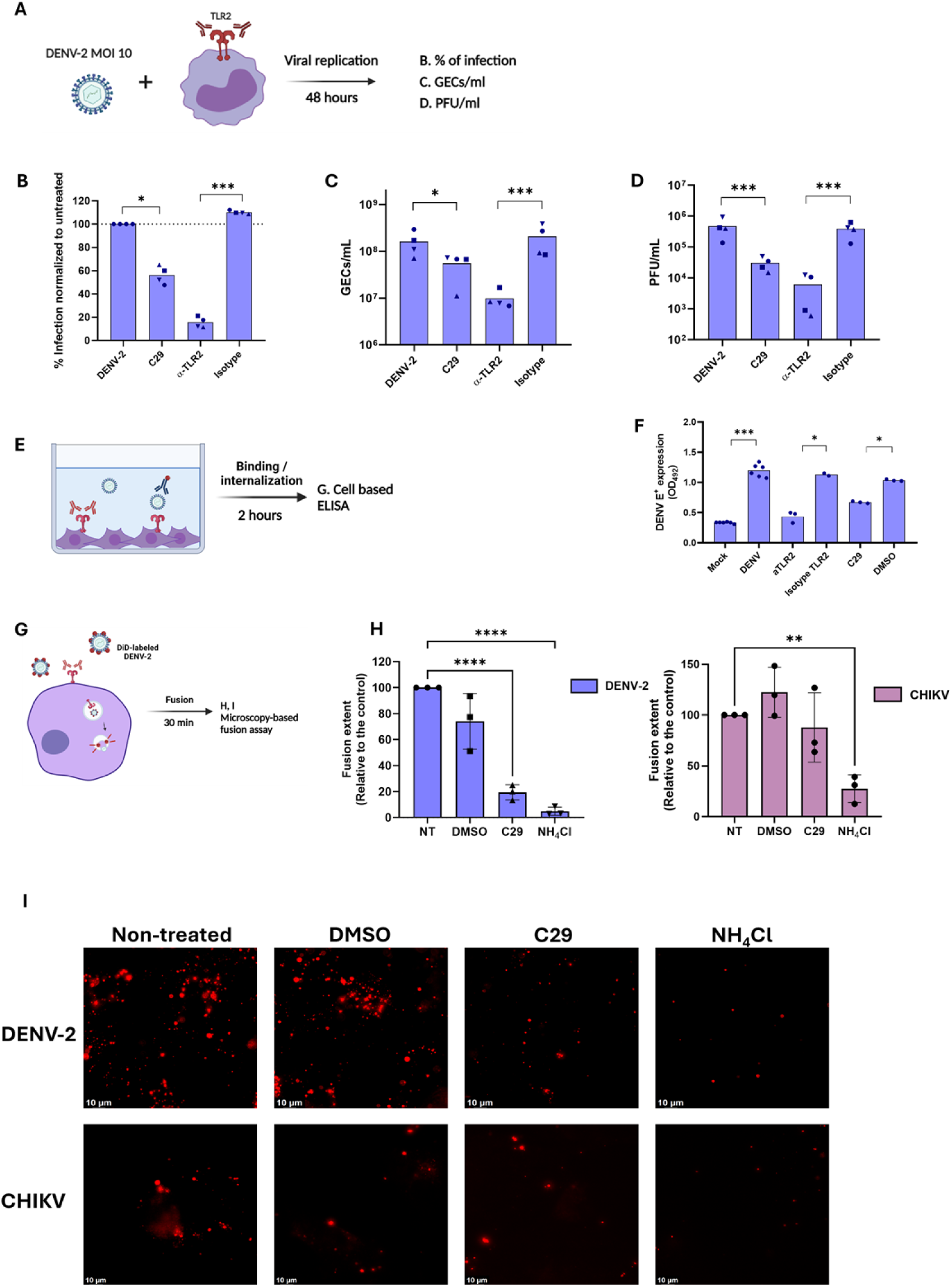
TLR2 activity supports early events required for productive DENV2 infection. (A) Experimental design (n= 4 different donors). (B) Percentage of DENV2 E-positive monocytes after treatment with C29, anti-TLR2, or corresponding controls. (C) Genome-equivalent copies GECs and (D) infectious titers (plaque-forming units, PFU) in supernatants at 48 hpi. Dashed line indicates the levels of residual inoculum (GECs and PFUs) after its removal at 2hpi. (E) Design of the cell-based virus-association assay. (F) Cell-associated E-protein signal in HEK-Blue hTLR2 cells after 2 h. (G) Quantification of DiD dequenching, (H) percentage inhibition, and (I) representative images after treatment with C29, DMSO, or NH4Cl. Data represent independent donors or experiments as indicated. Statistical tests and significance thresholds are shown in the figure. For statistical analysis Kruskal Wallis test and two-way ANOVA with a Dunnett post hoc test were used to determine significance with a 95% confidence interval (***p < 0.001, **p < 0.01, *p < 0.05).

Because these endpoint measurements could not distinguish an early entry-associated effect from a later effect on replication, we next examined events closer to virus uptake. In a cell-based assay using HEK-Blue hTLR2 cells, DENV2 particles associated with the cells during a 2-h exposure were detected with an antibody against the viral E protein (Fig. 1E). Treatment with either C29 or the TLR2-blocking antibody reduced the cell-associated E-protein signal compared with the corresponding vehicle or isotype control (Fig. 1F). We then used DiD-labelled DENV2 to assess membrane fusion. Dilution of DiD into the endosomal membrane after fusion causes fluorescence dequenching and provides a measure of the overall fusion signal. C29 treatment reduced this signal by approximately fourfold relative to untreated cells, while NH4Cl strongly inhibited fusion, as expected (Fig. 1G-I). To further assess the specificity of the C29 inhibitor, a TLR2-independent virus, chikungunya virus (CHIKV) was also included in the fusion assay. The same experimental procedure was followed using DiD-labelled CHIKV [32]. In contrast to DENV2, C29 treatment did not affect CHIKV fusion, indicating that the inhibitory effect of C29 is specific to TLR2-dependent DENV entry. Corresponding fluorescence images merged with brightfield/DIC images, providing cellular context for the observed fusion signals, are shown in Fig. S5.

These association and fusion experiments were performed in model cell lines and therefore do not establish direct binding of DENV2 to TLR2 or identify TLR2 as a bona fide entry receptor. Nevertheless, together with the primary-monocyte infection data, the results support the conclusion that TLR2-linked processes act early, at the level of virus-cell association, internalization, intracellular trafficking, and/or membrane fusion.

### TLR2-dependent infection enables inflammatory and antiviral responses that disrupt the endothelial barrier

Having established that TLR2 activity was required for efficient infection, we next asked whether this early dependence determined the inflammatory activity of infected monocytes. Monocytes were infected in the presence or absence of TLR2 blockade, and their cell-free supernatants were transferred to confluent HUVEC monolayers (Fig. 2A). Supernatants from DENV2-infected monocytes increased endothelial expression of E-selectin, ICAM-1, and VCAM-1. Blocking TLR2 at the cell surface or inhibiting its TIR domain with C29 markedly reduced this response (Fig. 2B). Direct exposure of HUVECs to DENV2 did not induce comparable activation (Fig. S6), indicating that endothelial activation was driven predominantly by soluble factors released from infected monocytes rather than by residual virus in the transferred supernatants.

**Fig. 2.**
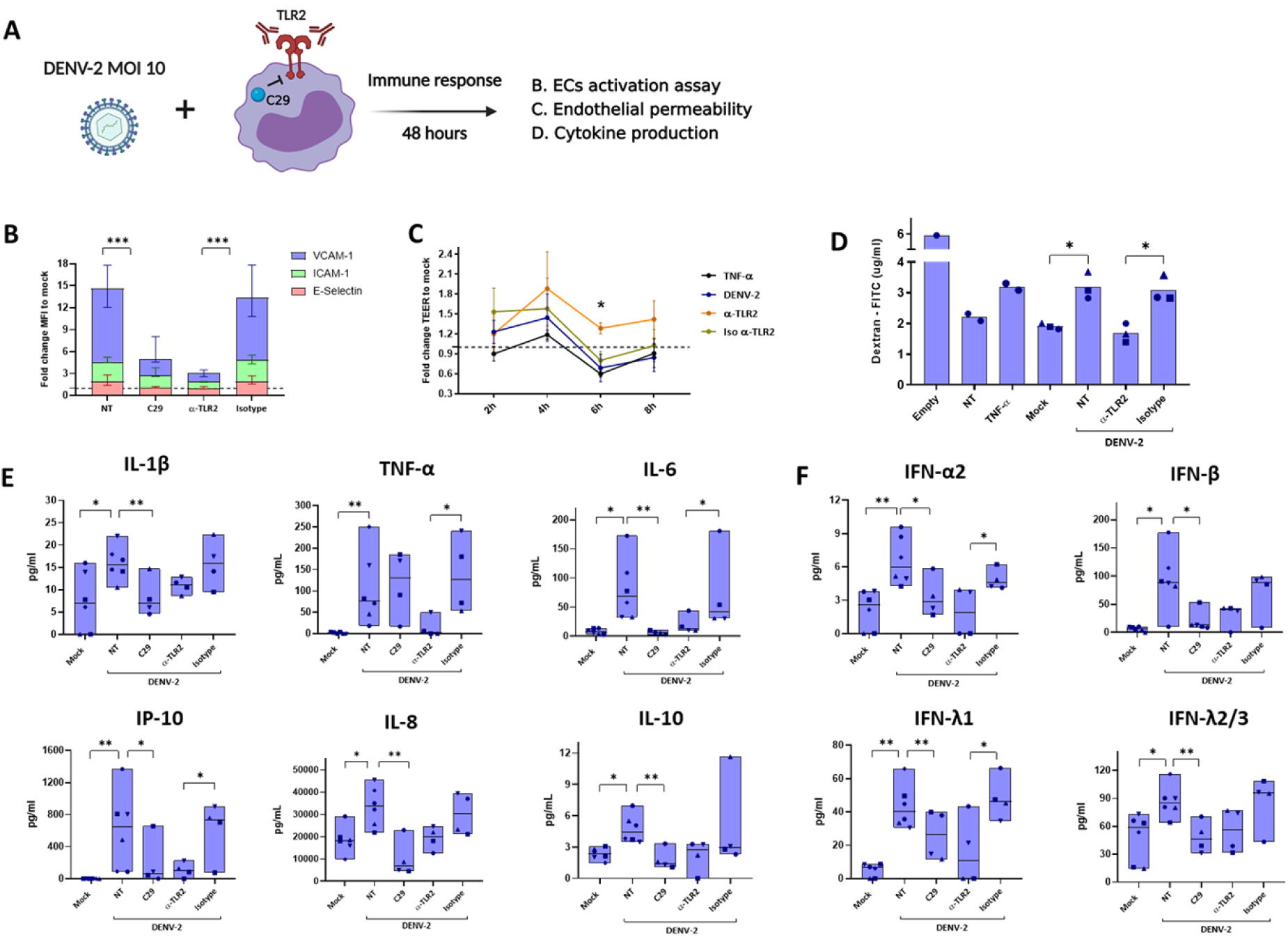
TLR2-dependent infection enables inflammatory and antiviral responses that impair endothelial barrier integrity. (A) Experimental design. (B) E-selectin, ICAM-1, and VCAM-1 expression in HUVECs exposed to supernatants from DENV2-infected monocytes treated with anti-TLR2, C29, or controls. (C) Relative TEER and (D) dextran-FITC flux across HUVEC monolayers. TNF-α served as a positive control. (E) Inflammatory mediators and (F) interferons in monocyte supernatants at 48 hpi. Data represent independent donors. Statistical tests and significance thresholds are shown in the figure. For statistical analysis two-way ANOVA with a Dunnett post hoc test was used to determine significance with a 95% confidence interval (***p < 0.001, **p < 0.01, *p < 0.05).

Endothelial activation, however, does not necessarily imply loss of barrier function. We therefore measured transendothelial electrical resistance (TEER) and dextran-FITC flux across HUVEC monolayers. Supernatants from DENV2-infected monocytes caused a transient but significant reduction in TEER and increased dextran passage, with effects comparable to those induced by the positive control TNF-α (Fig. 2C,D). Both changes were prevented when the monocytes had been infected under TLR2-blocking conditions. These findings show that the consequences of TLR2-dependent infection extend beyond induction of endothelial adhesion molecules and include measurable impairment of endothelial barrier integrity.

To define the soluble response associated with this phenotype, we quantified inflammatory and antiviral mediators in monocyte supernatants. DENV2 infection increased IL-1β, TNF-α, IL-6, IL-8, IP-10, IL-10, IFN-α2, IFN-β, IFN-λ1, and IFN-λ2/3, whereas IL-12p70, GM-CSF, and IFN-γ remained below the limit of detection (Fig. 2E,F). TLR2 inhibition reduced most of the DENV-induced mediators, in agreement with the endothelial activation and permeability results. Taken together, these data place TLR2-dependent infection upstream of a broad inflammatory and antiviral response capable of disrupting endothelial function.

### CD14 and CD36 contribute to infection and inflammatory output in purified monocytes

To investigate whether the receptor context of TLR2 influences DENV2 infection, we first compared basal expression of TLR2 and associated receptors, including CD14, CD36, TLR1, TLR6, and TLR10, among monocyte subsets (gating strategy in Fig. S7). TLR2 expression was comparable across CM, IM, and NM (Fig. 3A). CD14, CD36, TLR1, and TLR6 were more abundant on CM and IM than on NM, whereas TLR10 expression was higher in IM and NM.

**Fig. 3.**
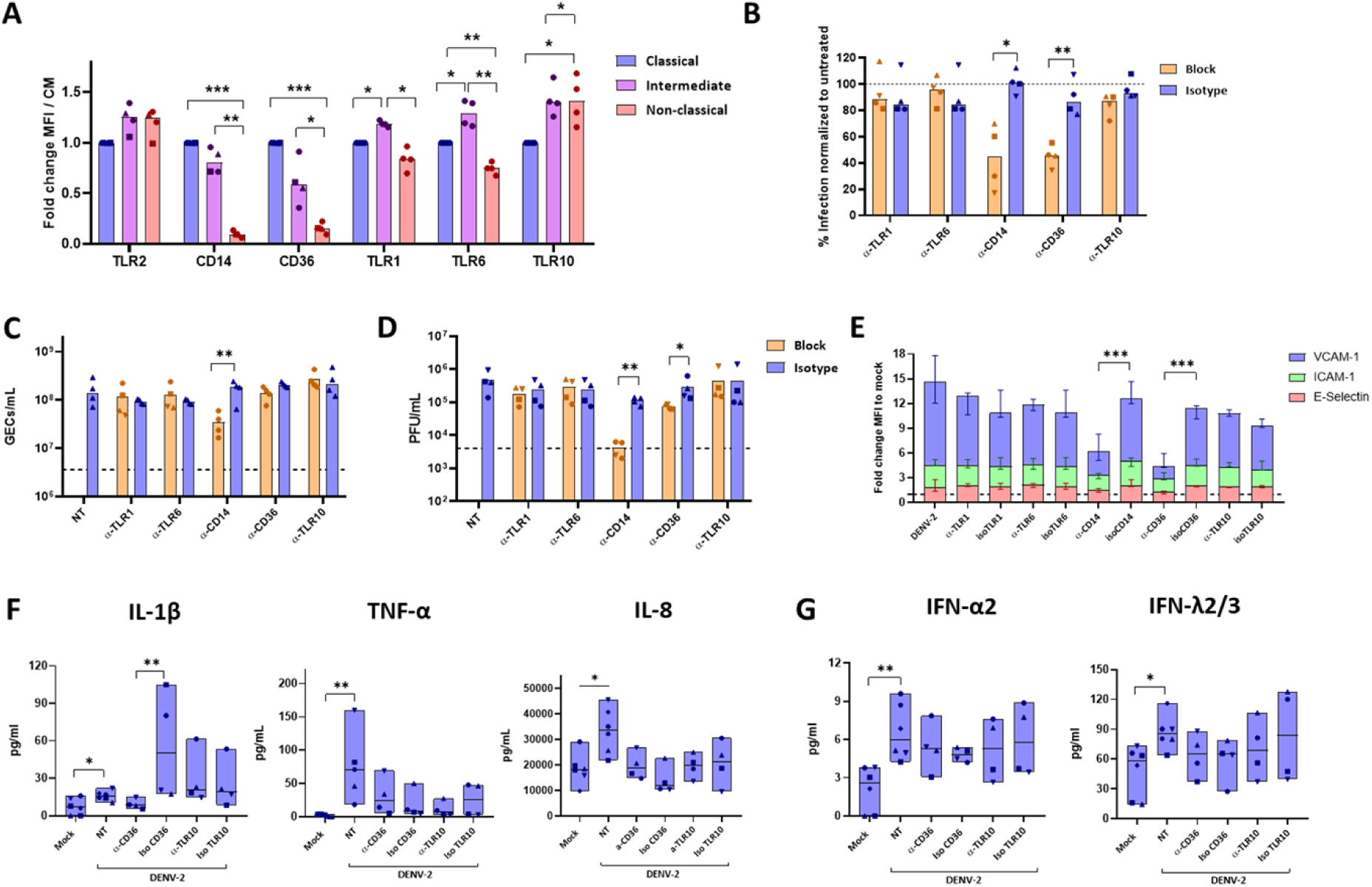
CD14 and CD36 contribute to DENV2 infection and inflammatory output in purified monocytes. (A) Expression of TLR2, CD14, CD36, TLR1, TLR6, and TLR10 in classical, intermediate, and non-classical monocytes. (B) Percentage of E-positive monocytes, (C) GECs and (D) infectious titers after blockade of TLR1, TLR6, TLR10, CD14, or CD36. (E) Endothelial activation is induced by monocyte supernatants. (F, G) Cytokine concentrations. Data represent independent donors. Statistical tests and significance thresholds are shown in the figure. For statistical analysis two-way ANOVA with a Dunnett post hoc test was used to determine significance with a 95% confidence interval (***p < 0.001, **p < 0.01, *p < 0.05).

Before testing the contribution of individual receptors, we confirmed that the blocking conditions were non-cytotoxic and functionally active using PAM2- and PAM3-induced responses (Fig. S8). Blocking TLR1, TLR6, or TLR10 did not alter the frequency of DENV2-infected monocytes, GEC production, infectious-virus output, endothelial activation, or cytokine production (Fig. 3B-G). These canonical TLR2 partners were therefore dispensable under the conditions tested. By contrast, CD14 blockade reduced the infected cell mass, GEC production, infectious-virus yield, and endothelial-activating activity. CD36 blockade produced a more modest phenotype: it reduced infected-cell mass and infectious-virus yield, but did not significantly reduce GEC production, and its effects on individual cytokines were limited. These findings identify CD14 and, to a lesser extent, CD36 as contributors to DENV2 infection in purified monocytes.

The receptor-blocking results require cautious interpretation. In our recent PBMC study, CD14 blockade reduced the staining accessibility of TLR2 and CD32 [23]. Therefore, reduced infection after CD14 blockade may reflect altered accessibility or organization of a receptor complex rather than an independent entry function of CD14. Moreover, CD36 was dispensable in the PBMC system [23], suggesting that its contribution is conditional on cellular context or experimental conditions.

### Classical monocytes selectively depend on the TLR2 axis for productive infection

The clinical association between sustained TLR2 expression on classical monocytes and progression to severe dengue prompted us to determine whether monocyte populations depend differently on the TLR2 axis [20]. Ideally, classical, intermediate, and non-classical monocytes would have been tested separately. However, the low frequencies of intermediate and non-classical monocytes did not allow recovery of sufficient cell numbers for parallel infection experiments. We therefore sorted CD16^−^ classical monocytes and a pooled CD16^+^ population comprising intermediate and non-classical monocytes (Fig. 4A; Fig. S9).

**Fig. 4.**
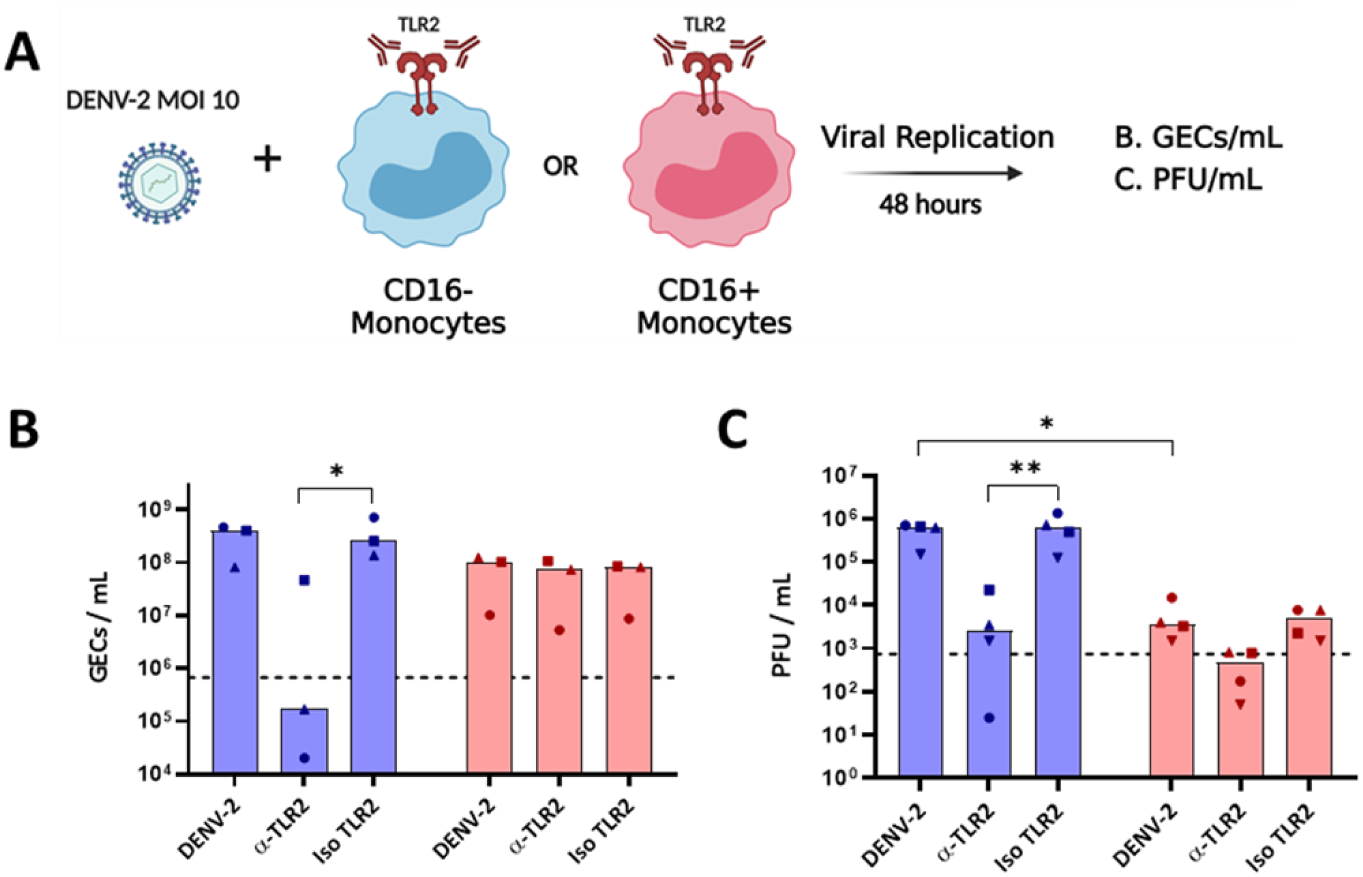
Productive infection of classical monocytes selectively depends on the TLR2 axis. (A) Sorting and infection strategy. (B) GECs and (C) infectious titers in supernatants of sorted CD16^−^ classical and pooled CD16^+^ intermediate/non-classical monocytes infected in the presence of anti-TLR2 or isotype control. Dashed lines indicate residual inoculum or the assay detection limit as defined in the figure. Data represent independent donors. Statistical tests and significance thresholds are shown in the figure. For statistical analysis two-way ANOVA with a Dunnett post hoc test was used to determine significance with a 95% confidence interval (**p < 0.01, *p < 0.05).

In the absence of TLR2 blockade, CD16^−^ and pooled CD16^+^ monocytes produced broadly comparable concentrations of viral genomes, with median values of 4.1 × 10^8^ and 1.1 × 10^8^ GECs/mL, respectively (Fig. 4B). The production of infectious virus differed more clearly. CD16^−^ monocytes yielded a median of 6.3 × 10^5^ PFU/mL, whereas pooled CD16^+^ monocytes produced a median of 3.6 × 10^3^ PFU/mL (Fig. 4C). This divergence between viral genomes and infectious progeny suggests that the populations differ not only in susceptibility but also in the efficiency with which infection progresses to production of infectious particles.

TLR2 blockade revealed a striking population-specific difference. Blocking TLR2 significantly reduced viral genome and infectious-particle production in CD16^−^ monocytes, but had no detectable effect in pooled CD16+ monocytes (Fig. 4B,C). Moreover, TLR2 blockade lowered infectious-virus production by CD16^−^ cells to approximately the level observed in untreated CD16^+^ cells. These findings identify classical monocytes as the population that most clearly relies on the TLR2 axis for productive infection. They do not establish which alternative route is used by intermediate or non-classical monocytes, because these subsets were not examined separately.

### Subset-specific TLR2 dependence determines endothelial-disruptive inflammatory output

We next asked whether the population difference in TLR2-dependent infection was reflected in the biological activity of the monocyte response. Supernatants from both DENV2-infected CD16^−^ and pooled CD16^+^ monocytes reduced endothelial barrier function, but the effect was stronger for CD16^−^ cells (Fig. 5A,B). Importantly, TLR2 blockade prevented the loss of endothelial integrity caused by CD16− monocyte supernatants but did not protect against the effect of CD16^+^ supernatants. Thus, the population in which productive infection depended on TLR2 was also the population in which TLR2 blockade prevented endothelial dysfunction.

**Fig. 5.**
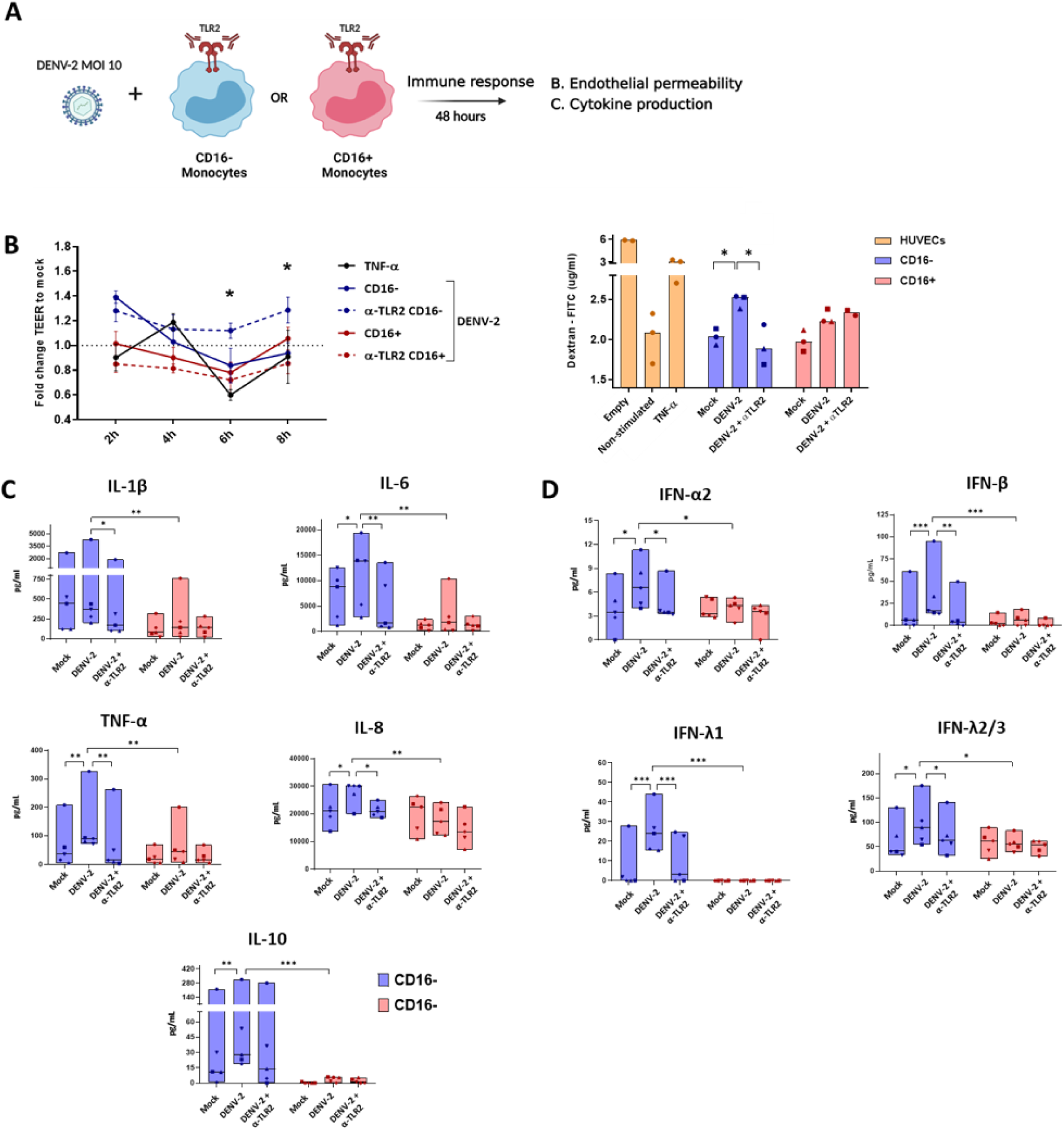
Subset-specific TLR2 dependence determines endothelial-disruptive inflammatory output. (A) Experimental design. (B) Relative TEER in HUVEC monolayers exposed to supernatants from infected CD16− or pooled CD16+ monocytes treated with anti-TLR2 or isotype control. (C) Inflammatory mediators and (D) interferons in monocyte supernatants. Data represent independent donors. Statistical tests and significance thresholds are shown in the figure. For statistical analysis two-way ANOVA with a Dunnett post hoc test was used to determine significance with a 95% confidence interval (***p < 0.001, **p < 0.01, *p < 0.05).

The cytokine profiles followed the same pattern. DENV2-infected CD16^−^ monocytes produced higher concentrations of IL-1β, IL-6, TNF-α, IL-8, IL-10, IFN-α2, IFN-β, IFN-λ1, and IFN-λ2/3 than pooled CD16^+^ monocytes (Fig. 5C,D). TLR2 blockade reduced these mediators in CD16^−^ cells but had no detectable effect in CD16^+^ cells. The lower DENV2-induced response of CD16^+^ monocytes did not reflect a general inability to respond through TLR2, because PAM3 stimulation elicited cytokine production in this population (Fig. S10). Together, these observations link the magnitude of the inflammatory response to population-specific use of the TLR2-associated infection route rather than to TLR2 signaling competence alone.

### NF-κB contributes to post-entry inflammatory and antiviral responses

The preceding experiments placed TLR2 activity upstream of both productive infection and cytokine release. We therefore examined NF-κB to determine how the inflammatory response was generated after infection had been established. DENV2 infection increased phosphorylation of the NF-κB p65 subunit, and TLR2 blockade reduced this response (Fig. 6A). Bay 11-7082 did not substantially affect monocyte viability and inhibited PAM3-induced TNF-α expression and endothelial-activating activity, confirming effective NF-κB inhibition (Fig. S11).

**Fig. 6.**
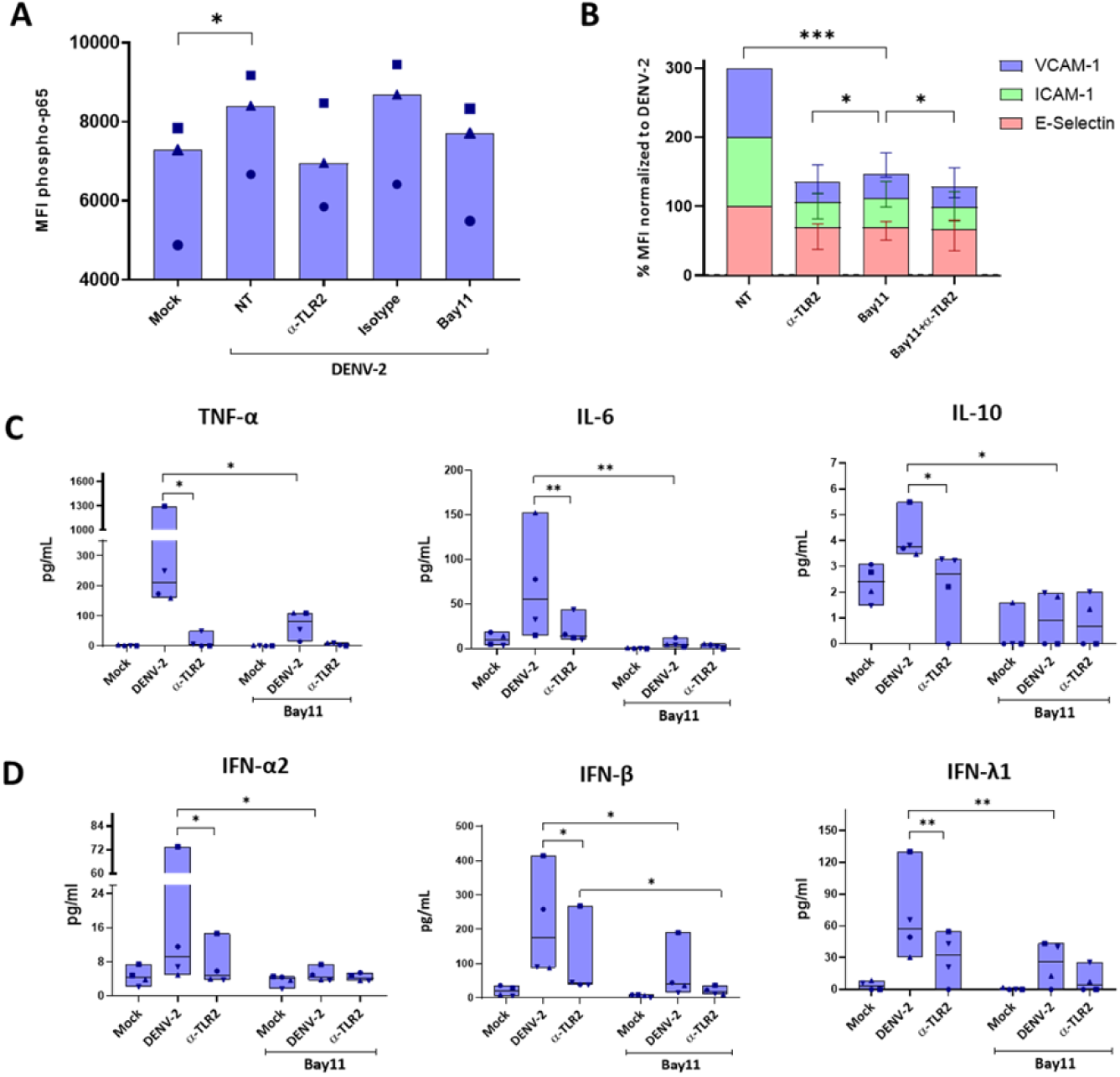
NF-κB contributes to post-entry inflammatory and antiviral responses. (A) Phosphorylated p65 in DENV2-infected monocytes treated with anti-TLR2, Bay 11-7082, or controls. (B) Endothelial activation induced by monocyte supernatants. (C) Inflammatory mediators and (D) interferons after treatment with anti-TLR2, Bay 11-7082, or both. Bay 11-7082 did not alter viral replication. Data represent independent donors. Statistical tests and significance thresholds are shown in the figure. For statistical analysis, two-way ANOVA with a Dunnett post hoc test was used to determine significance with a 95% confidence interval (***p < 0.001, **p < 0.01, *p < 0.05).

Bay 11-7082 reduced endothelial activation induced by DENV-infected monocyte supernatants and decreased production of TNF-α, IL-6, IL-10, IFN-α2, IFN-β, and IFN-λ1 (Fig. 6B-D). Combined treatment with Bay 11-7082 and the TLR2-blocking antibody produced a greater inhibitory effect than Bay 11-7082 alone for selected readouts. Because a formal interaction analysis was not performed, this result should not be interpreted as evidence of pharmacological synergy. Bay 11-7082 did not alter DENV2 replication (Fig. S12), separating the contribution of NF-κB to cytokine production from the earlier TLR2-associated effect on infection. These data position NF-κB as a major contributor to inflammatory and antiviral output downstream of infection.

### TLR3, but not TLR8, contributes to RNA-sensor-dependent responses in purified monocytes

Our recent PBMC study showed that intracellular RNA sensors contribute to the response after TLR2-supported direct or antibody-enhanced infection [23]. We therefore asked which RNA-sensing pathways were active in purified monocytes during direct infection. We focused on TLR3, which recognizes double-stranded RNA, and TLR8, which recognizes single-stranded RNA. Non-cytotoxic concentrations and functional activity of the inhibitors were confirmed using poly(I:C) and ssRNA40, respectively (Fig. S13).

TLR3 inhibition reduced the ability of DENV2-infected monocyte supernatants to activate endothelial cells and lowered IL-1β, IL-8, IFN-α2, IFN-β, and IFN-λ1 production (Fig. 7A, C, D). Combining TLR3 inhibition with TLR2 blockade further reduced endothelial activation and selected cytokines. Importantly, TLR3 inhibition did not alter the frequency of infected cells or production of infectious virus (Fig. S12). TLR3 therefore contributed to the response after infection without measurably controlling infection efficiency.

**Fig. 7.**
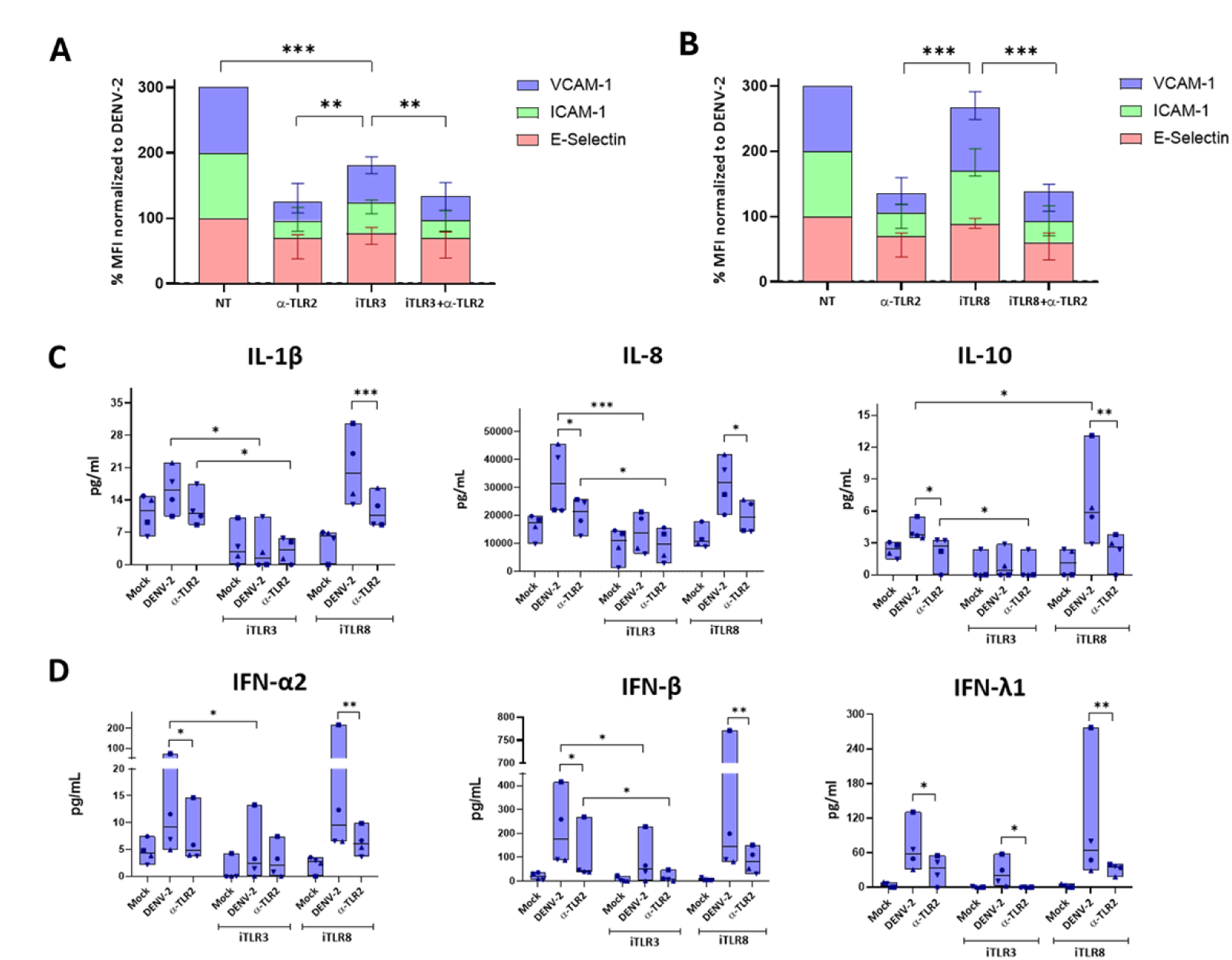
TLR3 contributes to inflammatory and antiviral responses in purified monocytes. (A,B) Endothelial activation induced by supernatants from monocytes treated with anti-TLR2, the TLR3 inhibitor, the TLR8 inhibitor, or combinations thereof. (C) Inflammatory mediators and (D) interferons in monocyte supernatants. TLR3 and TLR8 inhibition did not alter viral replication. Data represent independent donors. Statistical tests and significance thresholds are shown in the figure. For statistical analysis two-way ANOVA with a Dunnett post hoc test was used to determine significance with a 95% confidence interval (***p < 0.001, **p < 0.01, *p < 0.05).

In contrast, TLR8 inhibition did not significantly reduce endothelial activation or most cytokines, although IL-10 increased modestly (Fig. 7B-D). TLR8 inhibition also did not alter infection (Fig. S12). Thus, in purified monocytes undergoing direct infection, TLR3 contributed to inflammatory and antiviral output, whereas TLR8 was dispensable under the conditions tested. This differs from the mixed PBMC model, in which both TLR3 and TLR8 contributed, particularly during antibody-enhanced infection [23]. The difference suggests that RNA-sensor usage depends on cellular composition, uptake route, and infection context.

## DISCUSSION

Our understanding of TLR2 in dengue virus (DENV) infection has evolved through three complementary studies. We first linked TLR2-dependent responses in peripheral blood mononuclear cells (PBMCs) to NF-κB activation, inflammatory mediator production, endothelial activation, and progression to severe dengue [20]. We subsequently showed that TLR2 acts mainly during an early stage of direct and antibody-enhanced infection, whereas TLR3 and TLR8 shape downstream inflammatory and antiviral responses [23]. Here, we add another layer to this model by showing that CD16^−^ classical monocytes depend most strongly on the TLR2 axis. TLR2 blockade reduced productive infection, cytokine release, and endothelial-disruptive activity in classical monocytes, but not in pooled CD16^+^ monocytes. These findings show that TLR2 expression alone does not determine the outcome. Instead, monocyte identity determines whether cells couple TLR2-associated processes to productive infection and inflammatory output.

The early role of TLR2 is especially relevant because entry factors described for dendritic cells and differentiated macrophages do not fully explain why circulating monocytes support DENV infection. DENV can use DC-SIGN, heparan sulfate, and the mannose receptor in other myeloid cells, whereas circulating monocytes express little or no mannose receptor at the cell surface [13,14,34]. Earlier work also implicated a CD14-associated process in DENV attachment and/or entry into primary monocytes and macrophages [15]. Our data provide a possible molecular context for these observations. Blocking either the extracellular or intracellular domain of TLR2 reduced infection and progeny production in primary monocytes. TLR2 inhibition also reduced cell-associated viral antigen and DiD dequenching in model cell lines. Together, these findings support a role for TLR2-linked processes in virus association, uptake, trafficking, and/or fusion. However, they do not demonstrate direct DENV-TLR2 binding or establish TLR2 as a bona fide entry receptor. We therefore consider TLR2 an entry-associated host factor or permissiveness determinant.

CD14 and CD36 may help organize or internalize a TLR2-associated membrane complex rather than directly recognize DENV. Both receptors can transfer ligands and promote internalization of TLR2-containing complexes [24,26,27]. In purified monocytes, CD14 blockade produced the strongest phenotype, whereas CD36 blockade had a more limited effect on infectious-virus output. However, blocking antibodies can change receptor accessibility or membrane organization. In our recent PBMC study, CD14 blockade altered the staining accessibility of TLR2 and CD32, while CD36 blockade did not reduce infection [23]. We therefore interpret the contributions of CD14 and CD36 as context-dependent until genetic or biochemical approaches confirm their roles.

The selective dependence of classical monocytes on the TLR2 axis provides an important insight into how monocyte subsets may differ during DENV infection. However, the detection of viral antigen in cells with an intermediate or non-classical phenotype does not necessarily indicate that these cells were infected while already belonging to those subsets. Human monocytes follow a developmental continuum in which classical monocytes can differentiate into intermediate monocytes and subsequently into non-classical monocytes [35]. An infected classical monocyte may therefore acquire CD16 between virus exposure and phenotypic analysis and later be identified as an antigen-positive intermediate or non-classical monocyte. Consequently, post-infection phenotyping alone cannot distinguish phenotypic transition from genuine differences in initial susceptibility or post-entry permissiveness. Prospective separation of monocyte subsets before infection is therefore needed to compare these properties directly. In the present study, limited recovery of intermediate and non-classical monocytes required us to analyze these populations together, preventing us from resolving their individual susceptibility. Nevertheless, presorted CD16^−^ classical monocytes produced more infectious virus and showed a clear dependence on TLR2, whereas the pooled CD16^+^ population did not. Similarly, reports of viral antigen in CD16^+^ monocytes during HIV-1 or Zika virus infection [36,37] should be interpreted in light of the timing of subset identification and the possibility of differentiation after infection. Together, these considerations highlight the value of prospective experimental designs when assessing monocyte-subset susceptibility and permissiveness.

The difference between viral genome equivalents and infectious-virus yield suggests that monocyte subsets differ at more than the entry step. CD16^−^ and pooled CD16^+^ cells produced broadly comparable amounts of viral RNA, but CD16^+^ cultures yielded far fewer infectious particles. Viral RNA measurements do not show whether particles can complete another infection cycle. The lower infectious yield in CD16^+^ cultures may therefore reflect abortive infection, defective or incompletely mature particles, or restriction during assembly or release. Direct comparisons of attachment, internalization, fusion, replication, maturation, and release in separately presorted classical, intermediate, and non-classical monocytes will be needed to identify the restricted step.

Our inflammatory data support a serial model in which TLR2-associated infection enables subsequent RNA sensing. TLR3 inhibition reduced endothelial activation and cytokine production without altering infection, which places TLR3 downstream of entry. TLR8 did not measurably contribute in purified monocytes, whereas both TLR8 and TLR3 contributed in our PBMC and antibody-enhancement model [23]. Uptake route, antibody context, and communication among PBMC populations may therefore determine which RNA sensors become engaged. NF-κB also acted downstream of infection: its inhibition reduced inflammatory and antiviral mediators without changing viral replication. This finding extends earlier work linking DENV-induced NF-κB activation to TNF-α, inducible nitric oxide synthase, and macrophage migration inhibitory factor [38,39], while leaving the contribution of cytosolic RNA sensors unresolved.

The endothelial assays connect this monocyte-intrinsic pathway to a functional feature of dengue pathogenesis. Multiple mechanisms contribute to vascular dysfunction. DENV NS1 can directly weaken endothelial barriers, while monocyte-derived cytokines and chemokines can also promote endothelial activation and permeability [40–42]. In our experiments, direct exposure of HUVECs to the viral inoculum did not reproduce the effects of supernatants from infected monocytes. Those supernatants induced endothelial adhesion molecules, reduced TEER, and increased dextran flux. TLR2 blockade during infection prevented the barrier disruption caused by classical-monocyte supernatants. Monocyte-derived mediators are unlikely to explain vascular leakage on their own, but our data identify a biologically active pathway that links subset-specific infection to endothelial dysfunction.

Several limitations should be considered when interpreting our findings. Our conclusions are based primarily on pharmacological inhibition and antibody-mediated receptor blockade, while the virus-association and fusion assays were performed in model cell lines rather than primary monocytes. In addition, intermediate and non-classical monocytes were analyzed as a pooled CD16+ population, and the experiments were restricted to DENV2 strain 16681. We also did not identify the viral determinant involved in engaging the TLR2 axis or establish how TLR3 gains access to viral RNA. Finally, although TLR2 inhibition prevented monocyte-derived soluble mediators from disrupting endothelial barrier integrity ex vivo, these findings cannot be directly extrapolated to reduced vascular leakage in vivo.

In conclusion, our findings refine the current model of DENV recognition by identifying a monocyte-subset-specific checkpoint upstream of RNA sensing. TLR2-associated processes preferentially support productive infection of classical monocytes, thereby enabling TLR3- and NF-κB-dependent inflammatory and antiviral responses that compromise endothelial barrier function. The model provides a possible mechanistic explanation for the association between sustained TLR2 expression on classical monocytes and severe dengue [20]. The dual reduction in inflammatory and antiviral mediators after TLR2 blockade also argues against assuming that broad systemic TLR2 inhibition would be beneficial. A more selective strategy would require identification of the viral determinant and membrane complex involved in TLR2-associated infection, with the aim of disrupting viral exploitation of the pathway without suppressing protective innate immunity.

## ACKNOWLEDGMENTS

Our gratitude goes to Johan Teunis for excellent support with flow cytometry assays and to Timara Kuiper and Josée Plantinga (Endothelial Biomedicine & Vascular Drug Targeting group, UMCG) for their technical assistance in the culture of HUVEC.

The study was supported by UMCG and by the Colombian Ministry of Science, Technology, Knowledge, and Innovation (Minciencias). Part of this work was performed at the UMCG Imaging and Microscopy Centre (UMIC) supported by UMCG.

## DATA AVAILABILITY

All relevant data are within the manuscript and its supplementary information files.

## SUPPORTING INFORMATION LEGENDS

Fig. S1. Monocyte depletion reduces DENV infection, progeny production, and endothelial-activating activity in PBMC cultures.

Fig. S2. Purity of enriched monocyte cultures and efficiency of monocyte depletion.

Fig. S3. TLR2-blocking conditions are non-cytotoxic and inhibit PAM3-induced responses.

Fig. S4. Representative flow cytometry plots of DENV E-protein staining in monocytes.

Fig. S5. Representative DiD-labelled DENV and CHIKV fusion images merged with corresponding DIC images.

Fig. S6. Direct DENV exposure does not induce HUVEC activation or increased permeability under the conditions tested.

Fig. S7. Gating strategy for TLR2-axis expression in monocyte subsets.

Fig. S8. Viability and pathway-control experiments for blockade of TLR1, TLR6, TLR10, CD14, and CD36.

Fig. S9. Sorting strategy and post-sort purity of CD16^−^ and pooled CD16^+^ monocytes.

Fig. S10. Cytokine production by sorted monocyte populations after PAM3 stimulation.

Fig. S11. Viability and PAM3-control experiments for Bay 11-7082.

Fig. S12. DENV infection and replication after inhibiting NF-κB, TLR3, or TLR8.

Fig. S13. Viability and pathway-control experiments for TLR3 and TLR8 inhibition.

## REFERENCES

1. Bhatt S, Gething PW, Brady OJ, Messina JP, Farlow AW, Moyes CL, et al. The global distribution and burden of dengue. Nature. 2013;496: 504–507. doi:10.1038/nature12060

2. Dengue and severe dengue. [cited 15 Sept 2026]. Available: https://www.who.int/health-topics/dengue-and-severe-dengue

3. Dengue guidelines, for diagnosis, treatment, prevention and control. [cited 15 Sept 2026]. Available: https://www.who.int/publications/i/item/9789241547871

4. Butthep P, Chunhakan S, Yoksan S, Tangnararatchakit K, Chuansumrit A. Alteration of cytokines and chemokines during febrile episodes associated with endothelial cell damage and plasma leakage in dengue hemorrhagic fever. Pediatr Infect Dis J. 2012;31: e232–238. doi:10.1097/INF.0b013e31826fd456

5. Basu A, Chaturvedi UC. Vascular endothelium: the battlefield of dengue viruses. FEMS Immunology & Medical Microbiology. 2008;53: 287–299. doi:10.1111/j.1574-695X.2008.00420.x

6. Muller DA, Young PR. The flavivirus NS1 protein: molecular and structural biology, immunology, role in pathogenesis and application as a diagnostic biomarker. Antiviral Res. 2013;98: 192–208. doi:10.1016/j.antiviral.2013.03.008

7. Urcuqui-Inchima S, Patiño C, Torres S, Haenni A-L, Díaz FJ. Chapter 1 - Recent Developments in Understanding Dengue Virus Replication. In: Maramorosch K, Shatkin AJ, Murphy FA, editors. Advances in Virus Research. Academic Press; 2010. pp. 1–39. doi:10.1016/B978-0-12-385034-8.00001-6

8. den Boon JA, Diaz A, Ahlquist P. Cytoplasmic Viral Replication Complexes. Cell Host & Microbe. 2010;8: 77–85. doi:10.1016/j.chom.2010.06.010

9. Welsch S, Miller S, Romero-Brey I, Merz A, Bleck CKE, Walther P, et al. Composition and Three-Dimensional Architecture of the Dengue Virus Replication and Assembly Sites. Cell Host & Microbe. 2009;5: 365–375. doi:10.1016/j.chom.2009.03.007

10. Durbin AP, Vargas MJ, Wanionek K, Hammond SN, Gordon A, Rocha C, et al. Phenotyping of peripheral blood mononuclear cells during acute dengue illness demonstrates infection and increased activation of monocytes in severe cases compared to classic dengue fever. Virology. 2008;376: 429–435. doi:10.1016/j.virol.2008.03.028

11. Kou Z, Quinn M, Chen H, Rodrigo WWSI, Rose RC, Schlesinger JJ, et al. Monocytes, but not T or B cells, are the principal target cells for dengue virus (DV) infection among human peripheral blood mononuclear cells. J Med Virol. 2008;80: 134–146. doi:10.1002/jmv.21051

12. Neves-Souza PC, Azeredo EL, Zagne SM, Valls-de-Souza R, Reis SR, Cerqueira DI, et al. Inducible nitric oxide synthase (iNOS) expression in monocytes during acute Dengue Fever in patients and during in vitro infection. BMC Infect Dis. 2005;5: 64. doi:10.1186/1471-2334-5-64

13. Cruz-Oliveira C, Freire JM, Conceição TM, Higa LM, Castanho MARB, Da Poian AT. Receptors and routes of dengue virus entry into the host cells. FEMS Microbiol Rev. 2015;39: 155–170. doi:10.1093/femsre/fuu004

14. Miller JL, deWet BJM, Martinez-Pomares L, Radcliffe CM, Dwek RA, Rudd PM, et al. The Mannose Receptor Mediates Dengue Virus Infection of Macrophages. PLOS Pathogens. 2008;4: e17. doi:10.1371/journal.ppat.0040017

15. Chen YC, Wang SY, King CC. Bacterial lipopolysaccharide inhibits dengue virus infection of primary human monocytes/macrophages by blockade of virus entry via a CD14-dependent mechanism. J Virol. 1999;73: 2650–2657. doi:10.1128/JVI.73.4.2650-2657.1999

16. Geissmann F, Jung S, Littman DR. Blood Monocytes Consist of Two Principal Subsets with Distinct Migratory Properties. Immunity. 2003;19: 71–82. doi:10.1016/S1074-7613(03)00174-2

17. Wong KL, Yeap WH, Tai JJY, Ong SM, Dang TM, Wong SC. The three human monocyte subsets: implications for health and disease. Immunol Res. 2012;53: 41–57. doi:10.1007/s12026-012-8297-3

18. Ziegler-Heitbrock HW. Definition of human blood monocytes. J Leukoc Biol. 2000;67: 603–606. doi:10.1002/jlb.67.5.603

19. Merah-Mourah F, Cohen SO, Charron D, Mooney N, Haziot A. Identification of Novel Human Monocyte Subsets and Evidence for Phenotypic Groups Defined by Interindividual Variations of Expression of Adhesion Molecules. Sci Rep. 2020;10: 4397. doi:10.1038/s41598-020-61022-1

20. Aguilar-Briseño JA, Upasani V, Ellen BM ter, Moser J, Pauzuolis M, Ruiz-Silva M, et al. TLR2 on blood monocytes senses dengue virus infection and its expression correlates with disease pathogenesis. Nat Commun. 2020;11: 3177. doi:10.1038/s41467-020-16849-7

21. Kwissa M, Nakaya HI, Onlamoon N, Wrammert J, Villinger F, Perng GC, et al. Dengue virus infection induces expansion of a CD14(+)CD16(+) monocyte population that stimulates plasmablast differentiation. Cell Host Microbe. 2014;16: 115–127. doi:10.1016/j.chom.2014.06.001

22. Naranjo-Gómez JS, Castillo JA, Rojas M, Restrepo BN, Diaz FJ, Velilla PA, et al. Different phenotypes of non-classical monocytes associated with systemic inflammation, endothelial alteration and hepatic compromise in patients with dengue. Immunology. 2019;156: 147–163. doi:10.1111/imm.13011

23. ter Ellen BM, Punekar M, Castillo JA, Valdés-López JF, Upasani V, Sann S, et al. The contributions of TLR2, TLR8 and TLR3 to direct and antibody-dependent enhancement of dengue virus serotype 2 infection. npj Viruses. 2026;4: 24. doi:10.1038/s44298-026-00190-9

24. Jimenez-Dalmaroni MJ, Xiao N, Corper AL, Verdino P, Ainge GD, Larsen DS, et al. Soluble CD36 Ectodomain Binds Negatively Charged Diacylglycerol Ligands and Acts as a Co-Receptor for TLR2. PLOS ONE. 2009;4: e7411. doi:10.1371/journal.pone.0007411

25. Hasan U, Chaffois C, Gaillard C, Saulnier V, Merck E, Tancredi S, et al. Human TLR10 is a functional receptor, expressed by B cells and plasmacytoid dendritic cells, which activates gene transcription through MyD88. J Immunol. 2005;174: 2942–2950. doi:10.4049/jimmunol.174.5.2942

26. Shamsul HM, Hasebe A, Iyori M, Ohtani M, Kiura K, Zhang D, et al. The Toll-like receptor 2 (TLR2) ligand FSL-1 is internalized via the clathrin-dependent endocytic pathway triggered by CD14 and CD36 but not by TLR2. Immunology. 2010;130: 262–272. doi:10.1111/j.1365-2567.2009.03232.x

27. Raby A-C, Holst B, Le Bouder E, Diaz C, Ferran E, Conraux L, et al. Targeting the TLR Co-Receptor CD14 with TLR2-Derived Peptides Modulates Immune Responses to Pathogens. Science Translational Medicine. 2013;5: 185ra64–185ra64. doi:10.1126/scitranslmed.3005544

28. Silva MR, Briseño JAA, Upasani V, Ende-Metselaar H van der, Smit JM, Rodenhuis-Zybert IA. Suppression of chikungunya virus replication and differential innate responses of human peripheral blood mononuclear cells during co-infection with dengue virus. PLOS Neglected Tropical Diseases. 2017;11: e0005712. doi:10.1371/journal.pntd.0005712

29. Diosa-Toro M, Echavarría-Consuegra L, Flipse J, Fernández GJ, Kluiver J, van den Berg A, et al. MicroRNA profiling of human primary macrophages exposed to dengue virus identifies miRNA-3614-5p as antiviral and regulator of ADAR1 expression. PLoS Negl Trop Dis. 2017;11: e0005981. doi:10.1371/journal.pntd.0005981

30. Sachs LA, Schnurr D, Yagi S, Lachowicz-Scroggins ME, Widdicombe JH. Quantitative real-time PCR for rhinovirus, and its use in determining the relationship between TCID50 and the number of viral particles. J Virol Methods. 2011;171: 212–218. doi:10.1016/j.jviromet.2010.10.027

31. Ayala-Nuñez NV, Wilschut J, Smit JM. Monitoring virus entry into living cells using DiD-labeled dengue virus particles. Methods. 2011;55: 137–143. doi:10.1016/j.ymeth.2011.07.009

32. Hoornweg TE, van Duijl-Richter MKS, Ayala Nuñez NV, Albulescu IC, van Hemert MJ, Smit JM. Dynamics of Chikungunya Virus Cell Entry Unraveled by Single-Virus Tracking in Living Cells. J Virol. 2016;90: 4745–4756. doi:10.1128/JVI.03184-15

33. Mistry P, Laird MHW, Schwarz RS, Greene S, Dyson T, Snyder GA, et al. Inhibition of TLR2 signaling by small molecule inhibitors targeting a pocket within the TLR2 TIR domain. Proc Natl Acad Sci U S A. 2015;112: 5455–5460. doi:10.1073/pnas.1422576112

34. Noorman F, Braat EAM, Barrett-Bergshoeff M, Barbé E, van Leeuwen A, Lindeman J, et al. Monoclonal antibodies against the human mannose receptor as a specific marker in flow cytometry and immunohistochemistry for macrophages. Journal of Leukocyte Biology. 1997;61: 63–72. doi:10.1002/jlb.61.1.63

35. Patel AA, Zhang Y, Fullerton JN, Boelen L, Rongvaux A, Maini AA, et al. The fate and lifespan of human monocyte subsets in steady state and systemic inflammation. J Exp Med. 2017;214: 1913–1923. doi:10.1084/jem.20170355

36. Ellery PJ, Tippett E, Chiu Y-L, Paukovics G, Cameron PU, Solomon A, et al. The CD16+ Monocyte Subset Is More Permissive to Infection and Preferentially Harbors HIV-1 In Vivo. J Immunol. 2007;178: 6581–6589. doi:10.4049/jimmunol.178.10.6581

37. Michlmayr D, Andrade P, Gonzalez K, Balmaseda A, Harris E. CD14+CD16+ monocytes are the main target of Zika virus infection in peripheral blood mononuclear cells in a paediatric study in Nicaragua. Nat Microbiol. 2017;2: 1462–1470. doi:10.1038/s41564-017-0035-0

38. Cheng Y-L, Lin Y-S, Chen C-L, Wan S-W, Ou Y-D, Yu C-Y, et al. Dengue Virus Infection Causes the Activation of Distinct NF-κB Pathways for Inducible Nitric Oxide Synthase and TNF-α Expression in RAW264.7 Cells. Mediators Inflamm. 2015;2015: 274025. doi:10.1155/2015/274025

39. Chen L-C, Shyu H-W, Lei H-Y, Chen S-H, Liu H-S, Lin Y-S, et al. Dengue Virus Infection Induced NF-κB-dependent Macrophage Migration Inhibitory Factor Production. American J of Infectious Diseases. 2008;4: 22–31. doi:10.3844/ajidsp.2008.22.31

40. Kelley JF, Kaufusi PH, Nerurkar VR. Dengue hemorrhagic fever-associated immunomediators induced via maturation of dengue virus nonstructural 4B protein in monocytes modulate endothelial cell adhesion molecules and human microvascular endothelial cells permeability. Virology. 2012;422: 326–337. doi:10.1016/j.virol.2011.10.030

41. Lee Y-R, Liu M-T, Lei H-Y, Liu C-C, Wu J-M, Tung Y-C, et al. MCP-1, a highly expressed chemokine in dengue haemorrhagic fever/dengue shock syndrome patients, may cause permeability change, possibly through reduced tight junctions of vascular endothelium cells. J Gen Virol. 2006;87: 3623–3630. doi:10.1099/vir.0.82093-0

42. Puerta-Guardo H, Glasner DR, Espinosa DA, Biering SB, Patana M, Ratnasiri K, et al. Flavivirus NS1 Triggers Tissue-Specific Vascular Endothelial Dysfunction Reflecting Disease Tropism. Cell Rep. 2019;26: 1598–1613.e8. doi:10.1016/j.celrep.2019.01.036

